# The Growth Inhibitory study of Methanol Extract and Fractions of Seeds of *Chrysobalanus icaco* (Chrysobalanceae)

**DOI:** 10.1101/2025.09.23.676451

**Authors:** Jacinta Enoh Apitikori-owumi, Mubo Adeola Sonibare, Nwankwo Lawrence Uchenna, Nweke Chifumnaya Solomon, Akpomadaye Efe Cletus, Firdous Sayed Mohammed

## Abstract

**Background:** *Chrysobalanus icaco* L. (Chrysobalanaceae) is a spice commonly used by the Ijaws, Itsekiris, and Urhobos in Delta State. In ethnomedicine, it helps treat conditions such as diarrhoea, diabetes, high cholesterol, infections, and inflammation.

**Objective:** The study evaluated the growth inhibitory effect of *C. icaco* seed and its chromatographic fractions.

**Method:** The powdered seed sample of *C. icaco* was extracted with 70% ethanol by cold maceration, and the extract was screened for the presence of secondary metabolites. The extract was subjected to a vacuum liquid chromatographic technique, and the extract and fractions were subjected to bench-top biological assay using the growth inhibitory radicle of Sorghum bicolor at a concentration (1-30 mg/mL) for duration of 96 h. One way analysis of variance (ANOVA) was used in data analysis and was represented as Mean Standard Error of Mean (SEM).

**Results:** Secondary metabolites present in the extract were alkaloids, tannins, flavonoids, phenolic compounds, reducing sugar, terpenoids, proteins and amino acids, and phytosterols. The results revealed that the length of seed radicles progressively increased over time. The results showed that the length of seed radicles progressively increased over time. The extract reduced the length of the seed radicle by 14.51-58.39% and 66.44-84.74% at 20 and 30 mg/mL. The bulked vlc fractions BAF, BBF, and BDF significantly reduced the length of the radicle at 5 and 10 mg/mL, and bulked vlc BCF was effective at 10 mg/mL only.

**Conclusion:** The study showed that the extract of *C. icaco* and its vlc fractions possess an anti-proliferative effect with medical relevance in the development of anticancer therapy. The growth inhibitory effects of the extract of *C. icaco* and its vlc fraction on *Sorghum bicolor* could be linked to the phytochemicals present, associated with potential anticancer activity.

## Introduction

Cancer is regarded as one of the major public health issues and causes of death all over the world [1]. It is predicted that this number will rise by 40% in the near future (2030), accounting for roughly 13.1 million fatalities [2, 3]. According to reports from the World Health Organization, there were roughly 20 million new cases of cancer reported globally in 2022; of these, about 10.5 million were in men and 9.5 million in women. This indicates that the incidence of cancer is rising in years where there are 9.7 million deaths from the disease [4, 5] Thus, research into effective cancer treatments is still ongoing [6]. Medicinal plants are crucial for the development of new drugs [7].

Historically, people have utilized therapeutic herbs, and it’s possible that this practice is where modern medicine got its start [8]. Herbs and spices have been employed historically for culinary flavoring purposes and as a significant source of natural products in pharmaceuticals [9].

Some natural compounds have been acknowledged as important pharmacological players in the creation of new cancer-treatment medications. Research into medicinal plants used in treating tumor related ailments has become necessary due to the emergence of various forms of cancer diseases and the huge side effects usually encountered by cancer patients when treated with methods such as chemotherapy, radiotherapy and chemically derived drugs [10].

*Chrysobalanus icaco* commonly known as cocoplum, abajerủ belongs to the family of Chrysobalanceae and it is native to both Africa and America [11]. Biologically, *C. icaco* is reported to possess antioxidant, anticancer, antimicrobial, antidiabetic, diuretic, and anti-inflammatory activities [12]. *Chrysobalanus icaco* is also traditionally associated with the treatment of severe diarrhoea, vagina discharge, malaria, and bleeding ([13, 14, 15].

Studies on cell proliferation are an essential first stage in determining a test substance’s potential toxicity in people, including that of plant extracts or physiologically active chemicals derived from plants. Although cytotoxicity may suggest potential utility as an anti-cancer drug, little toxicity is crucial for the successful development of a pharmaceutical product. Some natural compounds have been identified as important players in the pharmacology of novel medications for the treatment of cancer [16]. While the majority of herbs and spice such as ginger, cinnamon and garlic have been reported to possess antiproliferative qualities, no study has been carried out on the growth inhibitory effect of the seed of *C. icaco*. Thus, the aim of this study is to determine the growth inhibitory effect of the *C. icaco* seed extract and chromatographic fractions of *C. icaco* seeds.

## Materials and Methods

### Material

The apparatus used during this study were: Whatmann No 42 Filter paper (Cole-Parmer Instrument, company, China), Petri Dish (Chongqing New World Trading Co., Ltd, China), Hot Air Oven (Huanyi Industry, Limited, Hong Kong), Refrigerator (Daliang Shunde, Foshan City, Guangdong Province, China), Mortar and Pestle (Chibez, Nig. Ltd, Onitsha, Nigeria), Electronic Weighing Balance (WANT Balance Instrument Co., Ltd, China), TLC pre-coated aluminium plate (Merck and Sigma-Aldrich, England), Silica Gel GF254 BOC Sciences, USA), Sinta Glass, No. 4 (Hebel Sinta FRP Co., Ltd, China), Rotary Evaporator (Infitek Inc, China), Buchner Flask (WF Education Group, England and Wales), Vaccum Pump (HACH, Dubai), Conical flask(WF Education Group, England and Wales), Beakers (250 mL, 500 mL) (WF Education Group, England and Wales), Measuring cylinder (1000 mL, 100 mL, 10 mL), Test Tubes(WF Education Group, England and Wales), Test Tube Racks, Sample Bottle (WF Education Group, England and Wales), Funnel(WF Education Group, England and Wales).

The following chemicals, reagents and drugs were used during this research work: Methanol, Hydrochloric acid (JHD, N-hexane, Iodine, Hydrochloric Acid (Merck and Sigma-Aldrich, England), Sulphuric acid (JHD, Guangdong Schi-Tech Ltd, China), dichloromethane(Merck and Sigma-Aldrich, England), Ethyl Acetate (Merck and Sigma-Aldrich, England), Methanol(Merck and Sigma-Aldrich, England), Guinea Corn Seeds, Cotton wool, DMSO_4_ Distilled Water, Fehling Solution A and B, Dragendorff’s Reagent, Mayer’s Reagent and NaOH.

### Plant Material Collection and Authentication

*Chrysobalanus icaco* plant was checked was with http://www.theplantlist..org and was assessed via http://www.worldfloraonline.org/taxon/wfo-0000830291. Dried fruits of *Chrysobalanus icaco* were brought from Main Market Abraka, Delta State, Nigeria. The fruit sample identity was confirmed at the Department of Botany and Biotechnology, University of Benin, Benin City, Nigeria where voucher number UBH-P606 was assigned.

### Preparation and Extraction of Seed Material

The seed were processes in the laboratory at the Department of Pharmacognosy and Traditional Medicine, Faculty of Pharmacy, Delta State University, Abraka. The seeds were separated from their shell and dried for one week at a room temperature in the laboratory and further dried using hot air oven at 40 ° C. The dried seeds were subsequently crushed with mortar and pestle, and then grinded into powder form using an electronic weighing balance and stored in an air tight container prior to extraction.

### Extraction of Powdered Seed

About 1000 g of the powdered seed of *C. icaco* was macerated in 70% methanol in a ratio of 1:4. The mixture was allowed to stand for 72 hours with constant daily agitation. Thereafter the macerated sample was filtered and the resulting filtrate was concentrated using a rotary evaporator to form a paste maintained at 40 ° C. The final extract was weighed and the yield calculated. It was then stored in a refrigerator at a temperature of 4 ° C for further use.

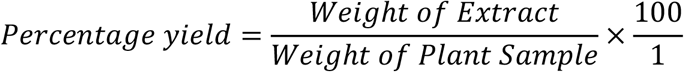

### Preliminary Phytochemical Screening

The preliminary phytochemical screening was conducted to detect the presence of secondary metabolites in the crude extracts and chromatographic fractions using method described previously [17].

### Vacuum Liquid Chromatography of *C. Icaco* Seed Crude Extract

Forty gram (40 g) of the methanol crude extract of C. *icaco* was prepared and packed on a Silica gel G with the size 30-70 µm in a Sinta Glass, No. 4 attached to a Buchner flask connected to a vaccum pump. The eluting solvents were 300 ml of hexane (100%), hexane – dichloromethane (75:25, 50:50, and 25: 75), dichloromethane (100%), dichloromethane ethyl acetate (75:25, 50:50, and 25:75), ethyl acetate (100%), ethyl acetate–methanol (75:25, 50:50, and 25:75) and methanol (100%). A total of thirteen (13) fractions were obtained. These fractions were then analyzed using Thin Layer Chromatographic techniques and similar fractions were bulked. A total of four (A1 (Fractions 3–7), A2 (fraction 8), A3 (Fractions 9–12 and A4 (fraction 13) fractions were obtained from the TLC analysis. The bulked chromatographic fractions these were further concentrated and subjected to biological assay against the germinating radicles of guinea corn using method as described below.

### Determination of the Anti-proliferative Effect of *C. Icaco* Seed Crude Extract

*Sorghum bicolor* (guinea corn), untreated was locally brought from Main Market Warri, Delta State, Nigeria. The corn seeds were cleansed according to the method described by Ikpefan[18] and Imade [19] with slight modification. Briefly, a handful of guinea corn seeds were placed in a beaker. 250 mL of 70% ethanol added to cleanse the seed in order to remove preservative. To the cleansed guinea corn in beaker, 250 mL of water was added and instantly decanted alongside the floating seeds. The seeds that were submerged were judged viable. The viable guinea corns were then allowed to air dry on filter paper before use. Twenty (20) guinea corn seeds were placed onto a Petri dish already underlined with filter paper and cotton wool and incubated in a dark cupboard for 96 h. The length of the radicle emerging from the seeds was then measured at 24, 48, 72 and 96 hours respectively. The guinea corn seeds used as control were treated with 10 mL of 2% DMSO in distilled water. The study was done in triplicates.

### Statistical Analysis

The data obtained were evaluated using GraphPad Prism 7.0. One way analysis of variance (ANOVA) was used in data analysis and was represented as Mean Standard Error of Mean (SEM).

## Results

### Percentage Yield of Extracts

The percentage yield of the crude extract of *C. icaco* is presented in Table 1. The result shows the yield of the extract and four (4) bulked chromatographic fractions of the crude extract of *C. icaco*. *C. icaco* Seed A2, A3 and A4 were 11.7%, 5%, 6% and 3.7% respectively.

**Table 1.** Percentage Yield of Crude Extract and Fractions of *C. icaco* Seed.

| **Sample** | **Weight (g)** | **Yield (%)** |
| --- | --- | --- |
| Crude extract | 105 | 10.5 |
| A1 | 3.5 | 11.7 |
| A2 | 1.5 | 5 |
| A3 | 1.8 | 6 |
| A4 | 1.1 | 3.7 |

### Phytochemical Analysis of Extract and Fractions

The preliminary phytochemical screening of the crude extract and chromatographic fractions of the leaf of *C. icaco* is presented in Table 2. The result showed the presence of twelve (12) metabolites which were saponins, tannins, phytosterol, flavonoids, terpenoids, alkaloids, steroids, reducing sugar, protein, carbohydrate, fat and oil and phlotannin in the crude extract. While a total of seven (7) metabolites were presents in the four bulked chromatographic fractions.

**Table 2.** Result of Preliminary Phytochemical Screening of Extract and Fractions.

| Phytochemical | Extract | A1 | A2 | A3 | A4 |
| --- | --- | --- | --- | --- | --- |
| Saponins | + | - | - | - | - |
| Tannins | + | + | + | + | + |
| Flavonoids | + | - | - | - | - |
| Cardiac glycosides | - | - | - | - | - |
| Terpenoids | + | + | + | + | + |
| Alkaloids | + | + | + | + | + |
| Steroids | + | - | - | - | - |
| Reducing sugar | + | + | + | + | + |
| Protein | + | + | + | + | + |
| Carbohydrate | + | + | + | + | + |
| Fat and oil | + | + | + | + | + |
| Phlotannin | + | - | - | - | - |

**Key:** + = present, - = absent

### Growth Inhibitory Effect of *C. icaco* Extract on Guinea Corn Radicle Length

A concentration-dependent radicle reduction was recorded. The average radicle length of 14.90±0.70 mm produced by the control seeds at 24 h was reduced 13.07±0.38, 12.63±0.91, 8.9±0.78, 6.20±0.85 and 5.00±1.53 by seeds treated with 1, 5, 10, 20 and 30 mg/mL of the crude extract. This decrease in radicle length was sustained up until the conclusion of the experiment (96 h) resulting in an average length of 38.36±1.60 mm against 31.10±1.61, 27.30±3.58, 25.43±0.61, 16.43±2.29 and 5.93±0 produced by seeds treated with 1, 5, 10, 20 and 30 mg/mL of the extract respectively showing 18.93, 28.83%, 33.71%, 57.17% and 84.54% (Table 3).

**Table 3:**
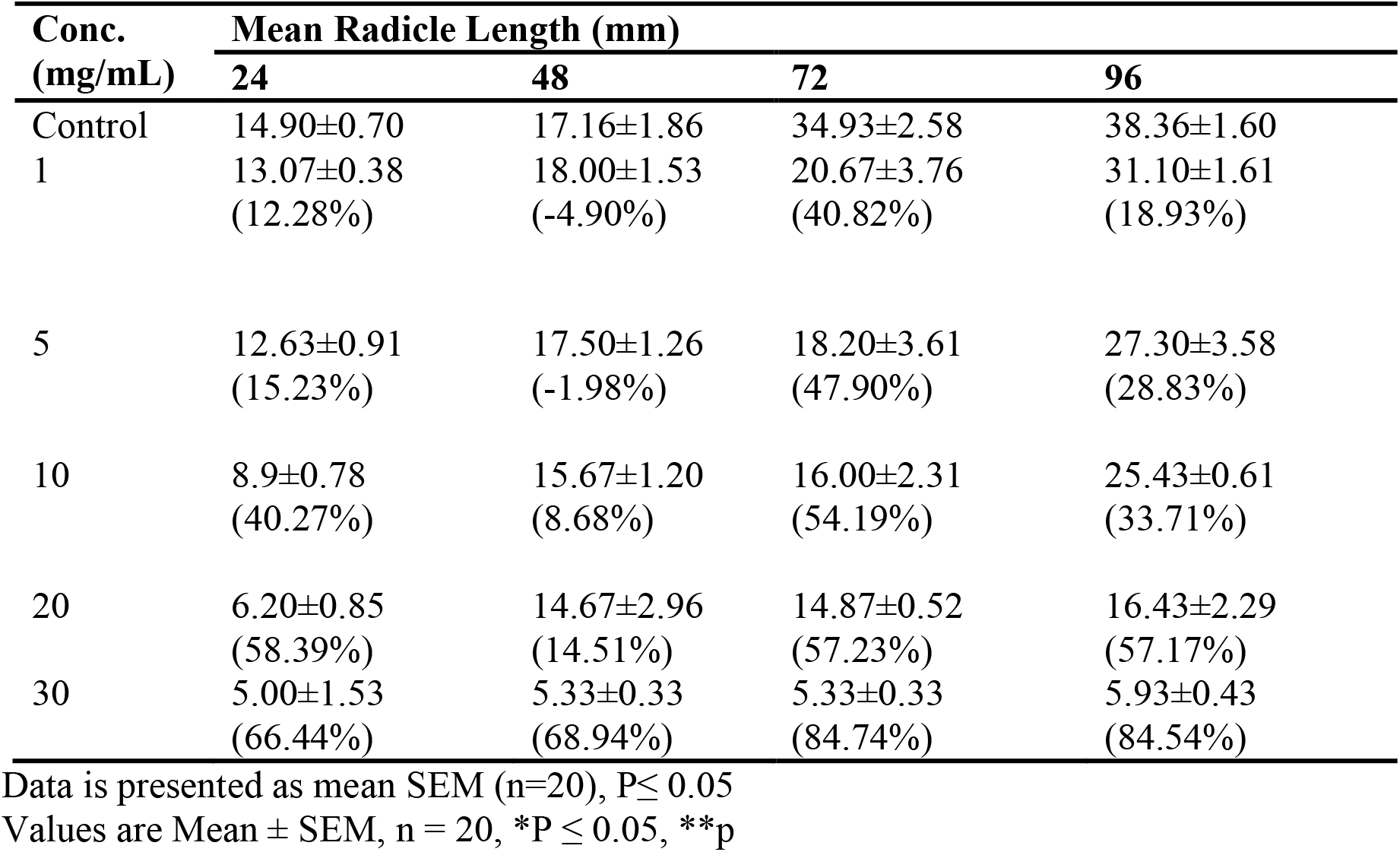
Growth Inhibitory Effect of *C. icaco* Extract on Guinea Corn Radicle Length.

| Conc.<br>(mg/mL) | Mean Radicle Length (mm) |  |  |  |
| --- | --- | --- | --- | --- |
|  | 24 | 48 | 72 | 96 |
| Control | 14.90±0.70 | 17.16±1.86 | 34.93±2.58 | 38.36±1.60 |
| 1 | 13.07±0.38<br>(12.28%) | 18.00±1.53<br>(-4.90%) | 20.67±3.76<br>(40.82%) | 31.10±1.61<br>(18.93%) |
| 5 | 12.63±0.91<br>(15.23%) | 17.50±1.26<br>(-1.98%) | 18.20±3.61<br>(47.90%) | 27.30±3.58<br>(28.83%) |
| 10 | 8.9±0.78<br>(40.27%) | 15.67±1.20<br>(8.68%) | 16.00±2.31<br>(54.19%) | 25.43±0.61<br>(33.71%) |
| 20 | 6.20±0.85<br>(58.39%) | 14.67±2.96<br>(14.51%) | 14.87±0.52<br>(57.23%) | 16.43±2.29<br>(57.17%) |
| 30 | 5.00±1.53<br>(66.44%) | 5.33±0.33<br>(68.94%) | 5.33±0.33<br>(84.74%) | 5.93±0.43<br>(84.54%) |

Data is presented as mean SEM (n=20), P≤ 0.05
Values are Mean ± SEM, n = 20, \*P ≤ 0.05, \*\*p

### Growth Inhibitory Effect of bulked Chromatographic fractions on Guinea Corn Radicle Length

As previously observed in the crude extract, the VLC bulked fractions were also seen to show a concentration-dependent outcome. At the end of 24 h, the control seeds produced a radicle length of 0.30±0.15mm which was reduced to 0.20±0.00 and 0.07±0.03 mm when treated with 5 and 10 mg/mL of fraction A1, This rise in radicle reduction was equally seem at 96 h when the control seeds’ 47.00±4.51 mm radicle length was decreased to 23.40±5.16 and 11.50±1.86 mm at comparable doses inferring 50.21% and 75.53% radicle length reductions (Table 4).

**Table 4:** Growth Inhibitory Effect of Bulk A1 on Guinea Corn Radicle Length.

| Concentration (mg/mL) | Time of incubation (hours) |  |  |  |
| --- | --- | --- | --- | --- |
|  | 24 | 48 | 72 | 96 |
| Control | $0.30 \pm 0.15$ | $15.17 \pm 3.71$ | $36.67 \pm 0.88$ | $47.00 \pm 4.51$ |
| 1 | $0.27 \pm 0.08$<br>(10.00%) | $3.87 \pm 1.67$<br>(74.49%) | $21.10 \pm 3.79$<br>(42.46%) | $23.40 \pm 5.16$<br>(50.21%) |
| 5 | $0.20 \pm 0.00$<br>(33.33%) | $1.00 \pm 0.70$<br>(93.41%) | $11.53 \pm 2.00$<br>(68.56%) | $11.50 \pm 1.86$<br>(75.53%) |
| 10 | $0.07 \pm 0.03$<br>(76.67%) | $0.23 \pm 0.09$<br>(98.48%) | $6.37 \pm 2.21$<br>(82.63%) | $7.40 \pm 1.78$<br>(84.26%) |

Data is presented as mean ± SEM (n=3)

Whereas the control seeds at 96 h gave 47.03±9.21 mm radicle length, seeds treated with fraction A2 at 5 and 10 mg/mL produce 7.20±1.10 mm and 2.13±0.52 mm radicle lengths respectively which denote 84.69% and 95.47% % reduction (Table 5). Still, at a comparable doses, bulked fraction A3 produced radicle lengths of 13.00±3.61 mm, 9.00±3.00 mm with an average radicle length of 26.50±4.81 mm produced by the control seeds (Table 6). While fraction A4 produced an average radicle length for the control seeds at 30.67±3.84 mm and at 5 mg/mL and 10 mg/mL gave 16.67±2.03 mm, 10.33±1.20 mm which suggest 45.65% and 66.32% (Table 7).

**Table 5:** Growth Inhibitory Effect of Bulk A2 on Guinea Corn Radicle Length.

| Concentration (mg/mL) | Time of incubation (hours) |  |  |  |
| --- | --- | --- | --- | --- |
|  | 24 | 48 | 72 | 96 |
| Control | $0.27 \pm 0.03$ | $32.13 \pm 6.44$ | $40.00 \pm 7.23$ | $47.03 \pm 9.21$ |
| 1 | $0.23 \pm 0.07$<br>(14.82%) | $12.13 \pm 6.40$<br>(62.25%) | $28.00 \pm 4.58$<br>(30.00%) | $16.77 \pm 7.03$<br>(64.34%) |
| 5 | $0.20 \pm 0.00$<br>(25.93%) | $2.90 \pm 1.72$<br>(90.97%) | $7.17 \pm 1.13$<br>(82.08%) | $7.20 \pm 1.10$<br>(84.69%) |
| 10 | $0.10 \pm 0.06$<br>(62.96%) | $0.43 \pm 0.29$<br>(98.66%) | $2.00 \pm 0.58$<br>(95.00%) | $2.13 \pm 0.52$<br>(95.47%) |

Data is presented as mean ± SEM (n=3)

**Table 6:**
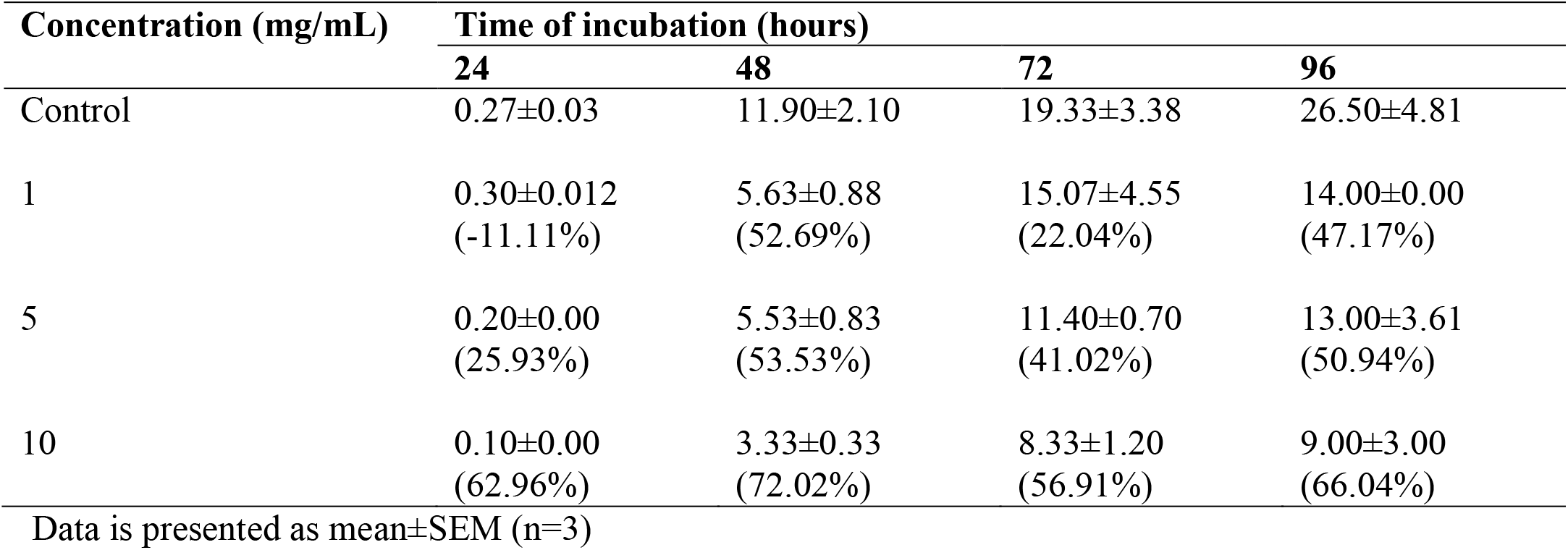
Growth Inhibitory Effect of Bulk A3 on Guinea Corn Radicle Length.

**Table 7:** Growth Inhibitory Effect of Bulk A4 on Guinea Corn Radicle Length.

| Concentration (mg/mL) | Time of incubation (hours) |  |  |  |
| --- | --- | --- | --- | --- |
|  | 24 | 48 | 72 | 96 |
| Control | 0.50±0.06 | 14.73±2.74 | 27.33±0.88 | 30.67±3.84 |
| 1 | 0.20±0.06<br>(60.00%) | 6.40±0.70<br>(56.55%) | 16.33±3.48<br>(40.25%) | 20.67±2.73<br>(32.61%) |
| 5 | 0.17±0.07<br>(66.00%) | 5.77±1.49<br>(60.83%) | 11.00±5.00<br>(59.75%) | 16.67±2.03<br>(45.65%) |
| 10 | 0.15±0.03<br>(70.00%) | 4.07±1.18<br>(72.37%) | 7.80±0.61<br>(71.46%) | 10.33±1.20<br>(66.32%) |

Data is presented as mean±SEM (n=3 \*P ≤ 0.05, \*\*p

## Discussion and Conclusion

Plant-based remedies have long been used to treat illnesses in people. Natural products are very significant source of new lead compounds for drug development research especially for infectious and cancer disease [20, 21].

Cancer, which can affect any region of the body is characterized by the fast and uncontrollable proliferation of aberrant cells that are resistant to planned cell killing [22]. Tumours that have the propensity to spread laterally may then result from this [23]. Cancer has emerged as one of the world’s most serious health issues in recent years, seriously affecting people’s life expectancy and quality of life. The International Agency for Research on Cancer (GLOBOCAN) database just released a report stating that 10 million people died and 19.3 million new cases of cancer were diagnosed in 2020 [24]. On the African continent, in particular in sub-Saharan Africa, this tendency is also noted. In fact, 520,348 cancer-related deaths and 801,702 new cases were reported in sub-Saharan Africa [24]. Because nearly every country in sub-Saharan Africa faces financial, economic, and health challenges, cancer treatment based on surgery, radiation, and chemotherapy is a significant barrier for these nations [25]. In an effort to overcome these challenges, scientists and researchers are looking more and more to medicinal plants as viable alternatives for the prevention and treatment of cancer. In fact, compared to traditional chemotherapeutic drugs, medicinal plants have bioactive chemicals that function as anticancer agents with fewer side effects and a more favourable toxicity profile [26].

To investigate the potential cytotoxic and growth-inhibitory effects of C. icaco seed extract, growth inhibitory effect, using guinea corn seeds (*Sorgum bicolor*), well-established, quick, and easy bioassay experimental models was used [18]. As previously mentioned, bioassays are a necessary component of natural product chemistry research projects [27]. Extracts have to be tested for biological activity, the active extracts chosen, fractionations guided by bioassays, and the bioactive components found and subsequently utilized. The fact that seed meristematic tissues tend to grow in suitable environments was a factor in the selection of guinea maize. While other plant seeds may have been utilized, Sorghum bicolor seeds were chosen due to their comparatively tiny size and capacity to provide a germination rate of approximately 90% in less than 24 hours [18].

Bioactive substances are essential for many different range of therapeutic applications, and some of them possess anticancer properties [28]. Preliminary phytochemical analysis of the extract was determined to know the class of phytochemical contained in it and the result showed the presence of saponins, tannins, phytosterol, flavonoids, terpenoids, alkaloids, steroids, reducing sugar, protein, carbohydrate, fat and oil and phlotannin. According to reports from previous research, varied phenolic compounds have been associated with cytotoxic and anti-proliferative properties against three melanocyte cell lines [29]. One possible explanation for the observed decrease in seed radicle length could be the presence of various kinds of secondary metabolites within the seed [18].

The study also examined the effect of *C. icaco* seed extract and chromatographic fractions on the growth inhibitory effect of guinea corn seed radicle. There was a notable concentration-dependent decrease in the radicle length of the Sorghum bicolor seeds when exposed to the methanol crude extract and fractions. It was noted that, in comparison to the control, the length of the radicle continuously decreased noticeably for the seeds as the incubation period increased. With a treatment dose of 30 mg/ml of the seed extract (5.33±0.33 mm), the result demonstrates a substantial growth inhibitory effect of the extract, with the largest growth reduction obtained at 96 h. The length of the seeds’ radicles was found to significantly decrease with increasing concentration in the methanol seed extract. The outcome also demonstrates that, with the exception of concentrations of 1, 5, and 10 mg/mL, the length of the radicle significantly decreased with an increase in incubation time.

The controls had an average of 38.36±1.60 mm while 20 mg/mL and 30 mg/mL of the crude extract had 16.43±2.29 mm and 5.93±0.43 mm respectively, showing 57.17% and 84.54% reduction in length respectively.

The bulked vlc fractions of A2 was seen to exhibit a significant inhibitory effect over bulked fractions A1, A3 and A4 has it has an average 7.20±1.10 mm at 5.0 mg/mL and 2.13±0.52 mm at 10 mg/mL after 96 h showing 84.69% and 95.47% with a reduction in length compared to 38.36±1.60 mm while fraction A3 had a significant reduction in the length of the guinea corn seed radicle at a concentration of 10 mg/mL. This may be explained by the fact that chemical constituents of the extract directly interfered with the biochemical processes of the guinea corn (*Sorghum bicolor*), causing the seed to proliferate at that time. The chromatographic fractions of the crude extract’s results showed that the fractions prevented radicle growth, and thus showing that, when isolated, might be just as powerful as the crude extract.

*Chrysobalanus icaco* seed extracts and its chromatographic fractions presented anti-proliferative properties against the radicles of guinea corn. The anti-proliferative properties may be attributed to the plant’s presence of some bioactive constituents, such as terpenoids, which may be connected to its use as an anticancer agent. To further support this assertion, additional research with cancer cell lines is recommended.

## Notes

### Competing Interest Statement

The authors have declared no competing interest.

